# Modeling enamel matrix secretion in mammalian teeth

**DOI:** 10.1101/525162

**Authors:** Teemu J. Häkkinen, S. Susanna Sova, Ian J. Corfe, Leo Tjäderhane, Antti Hannukainen, Jukka Jernvall

## Abstract

The most mineralized tissue of the mammalian body is tooth enamel. Especially in species with thick enamel, three-dimensional (3D) tomography data has shown that the distribution of enamel varies across the occlusal surface of the tooth crown. Differences in enamel thickness among species and within the tooth crown have been used to examine taxonomic affiliations, life history, and functional properties of teeth. Before becoming fully mineralized, enamel matrix is secreted on the top of a dentine template, and it remains to be explored how matrix thickness is spatially regulated. To provide a predictive framework to examine enamel distribution, we introduce a computational model of enamel matrix secretion that maps the dentine topography to the enamel surface topography. Starting from empirical enamel-dentine junctions, enamel matrix deposition is modeled as a diffusion-limited free boundary problem. Using laboratory microCT and synchrotron tomographic data of pig molars that have markedly different dentine and enamel surface topographies, we show how diffusion-limited matrix deposition accounts for both the process of matrix secretion and the final enamel distribution. Simulations reveal how concave and convex dentine features have distinct effects on enamel surface, thereby explaining why the enamel surface is not a straightforward extrapolation of the dentine template. Human and orangutan molar simulations show that even subtle variation in dentine topography can be mapped to the enamel surface features. Mechanistic models of extracellular matrix deposition can be used to predict occlusal morphologies of teeth.

**Author summary:** Teeth of most mammals are covered by a layer of highly mineralized enamel that cannot be replaced or repaired. The enamel layer is not uniform over the underlying dentine, and spatial regulation of enamel formation is critical for making a functional tooth. To explore which kind of mechanisms could underlie the complex patterns of enamel distribution, we present a computational model. Starting from tomography-imaged teeth from which enamel has been digitally removed, enamel is restored on dentine surfaces by simulating diffusion-limited secretion of enamel matrix. Our simulations show how the combination of subtle features of dentine and diffusion-limited secretion of the enamel matrix can substantially increase the complexity of the enamel surface. We propose that the strength of the diffusion-limited process is a key factor in determining enamel distribution among mammalian species.

## Introduction

Most mammalian species have their teeth covered by a layer of highly mineralized enamel. The thickness of the enamel layer relative to the tooth size ranges from thin to very thick. These differences among species, and also increasingly within the tooth crown, have been informative in studies focused on functional properties of teeth, taxonomy, and life history [1–9]. Even though mutations in genes required for enamel matrix secretion and maturation are known to affect the enamel thickness in mammals [10], relatively little is known about the regulatory changes that might underlie the variation in enamel thickness among populations or species [1, 11, 12]. Even less is known about the regulation of enamel thickness variation within the tooth crown, which contrasts with the increasing availability of 3D tomography data on various species. Analyses of such tomography data show that even though the enamel surface topography reflects the enamel-dentine junction (EDJ) topography, the enamel surface is not a simple extrapolation of the EDJ shape [13–15]. Because enamel distribution is not developmentally remodeled after formation, and because the internal structure of mineralized enamel retains developmental information, tomography data of fully formed teeth can be used to examine mechanisms underlying variation in enamel thickness. To provide mechanistic insights into the regulation of enamel thickness, here we combine tomography data on enamel distribution with a computational approach and introduce a model to simulate enamel matrix secretion.

## Results

### The model principles and simulation of artificial shapes

The enamel matrix is secreted by specialized epithelial cells, the ameloblasts. When the matrix secretion begins, ameloblasts detach from the EDJ and advance as a front secreting the enamel matrix that will later mineralize into the enamel (Fig 1A). The EDJ is defined by the mesenchymal dentine matrix, whose secretion begins first (Fig 1A). For empirical tests, we used EDJs of real teeth as the starting point to simulate matrix secretion. Matrix deposition is modeled as a diffusion-limited free boundary problem, motivated by the classical Stefan problem that models phase transition of undercooled liquid by assuming that the rate of phase transition from liquid to solid is limited by a diffusion process (see Methods and S1 Appendix) [16, 17]. Here we similarly assume that the growth of the matrix front is a diffusion-limited process: The advancement of the ameloblast layer is assumed to be limited by the diffusion of nutrients, by which we refer collectively to all the factors that ameloblasts require for the secretion of the matrix (Fig 1B). Biologically, both the extracellular milieu and capillary networks present in the enamel organ during matrix secretion [18, 19] can be hypothesized to limit the supply of nutrients in a diffusion-limited fashion. The model parameters adjust the nutrient diffusion rate, the amount of nutrients required for growth, and the interfacial tension or stiffness of the advancing ameloblast layer (Methods). Model equations are solved using the finite element method, and the matrix interface (the ameloblast layer) is tracked using the level set method (S1 Appendix). The source code of the Matlab implementation of the model is freely available (Methods). For computational efficiency, the model is implemented in 2D and 3D reconstructions are obtained by simulating multiple sections that are combined into volumes.

**Fig 1.**
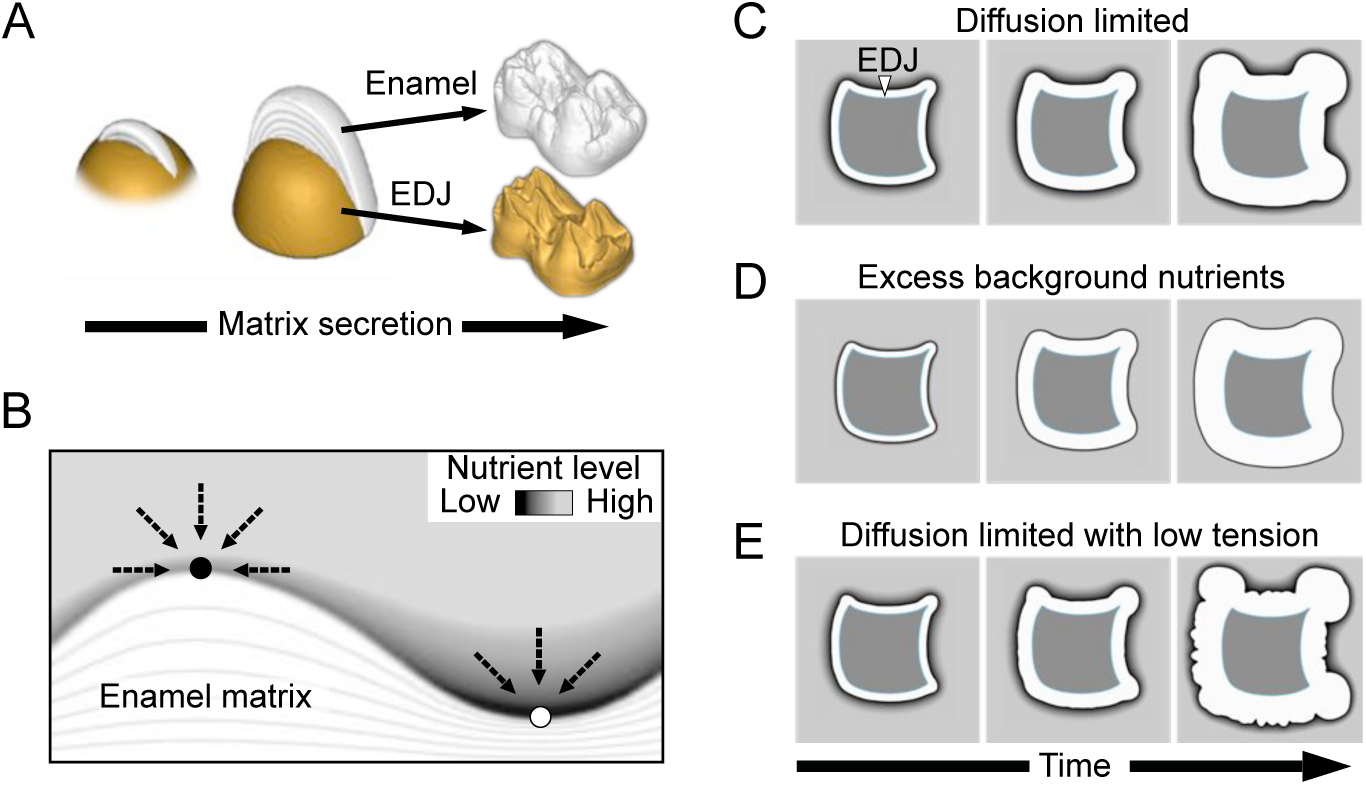
Modeling tooth enamel matrix secretion. (A) A schematic illustration of a cross section of enamel matrix being secreted on the top of dentine. In real teeth (a pig molar on the right), the enamel surface is not a linear representation of the dentine template (EDJ). (B) In a diffusion-limited model, differences in surface topography lead to ridges (black circle) receiving more nutrients (dashed arrows) than valleys (white circle). (C) Starting from a synthetic EDJ shape, diffusion-limited matrix deposition advances faster in convex than concave features. (D) Excess production of nutrients overcomes diffusion-limited effects and produces a uniform distribution of matrix. (E) Reducing interfacial tension of the simulation in (C) results in a crenulated matrix surface. For details of the model, see Methods, and parameters used in simulation are in S1 Table.

The fundamental component of the model is the assumption that the growth of the matrix requires a net influx of a diffusing nutrient substance. At the initial stage nutrients are present exterior to the dentine, which in real teeth, acts as an internal nutrient barrier [18]. The nutrients are further replenished over time by a constant background source exterior to the dentine (Methods). By controlling the relative amount of background production, we examine two hypothetical matrix secretion processes. The primary process tested is a diffusion-limited secretion in which concave surfaces are progressively exaggerated as the features protruding into the nutrient-rich domain receive more nutrients than the concavities (Fig 1C). An alternative process assumes excess availability of nutrients through strong background production, leading to a moving boundary of uniform thickness (Fig 1D). This latter process in fact closely approximates a simple geometric extrapolation of matrix thickness from the EDJ, which we use as a null hypothesis to demonstrate the non-linearity of the matrix deposition. Simulations of matrix secretion using a synthetic EDJ shape show that whereas convex EDJ surfaces result in relatively linear extrapolation of the enamel surface in both simulations (Fig 1C and 1D), concave surfaces of diffusion-limited simulations behave nonlinearly (Fig 1C). Additionally, reducing interfacial tension in the simulations increases small undulations in the moving front (Fig 1E), suggesting that lowered stiffness of the ameloblast layer may underlie crenulated enamel found in taxa such as *Chiropotes* (saki monkeys) with relatively smooth EDJ [14]. Next we applied the model to real teeth with convex and concave features (Fig 2A).

**Fig 2.**
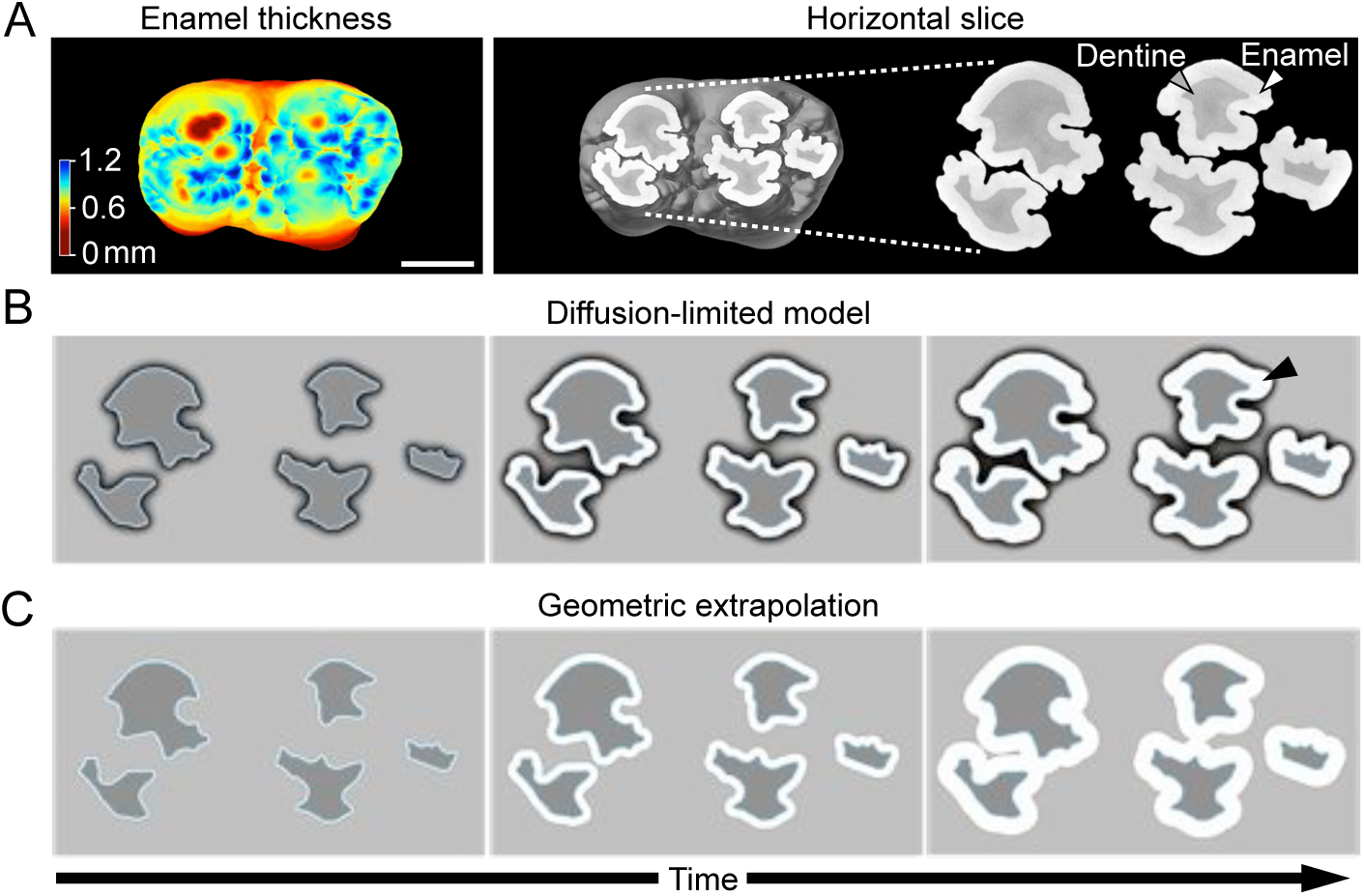
Diffusion-limited simulations approximate complex patterns of enamel thickness in the pig molar. (A) A heat map and a horizontal section of 3D tomography reconstruction of a pig molar shows the variable enamel thickness. (B) Using a horizontal EDJ section of a pig molar as an empirical template (A), diffusion-limited simulations of matrix secretion produce deep fissures present in concave surfaces. (C) Geometric extrapolation shows how the fissures are filled-in. In contrast, convex slopes are relatively similar between the simulations. Note how the enamel is redirected and remains separated between the buccal and lingual cusps in the real tooth (A) and diffusion-limited simulation (B). All images show occlusal views of the left lower first molar. Simulations run until the lateral matrix thicknesses approximate the empirical enamel thicknesses, see Methods, and for parameters S1 Table. Scale bar, 5 mm.

### Diffusion-limited simulations predict enamel distribution on pig molar teeth

To simulate enamel matrix secretion in real teeth, first we focused on domestic pig molar teeth which exhibit substantial variation in enamel thickness and EDJ topography (Fig 1A and 2A) [20]. EDJ and enamel surface shapes were reconstructed from microCT scans of first lower molars (Fig 2A, Methods). From the data, horizontal slices of cusps were extracted (Fig 2A) and the EDJs were used as the starting point for the simulations. The horizontal plane represents a relatively synchronous front of enamel matrix secretion and captures the complex EDJ morphology of the pig molars [20]. The simulations show that the diffusion-limited process reproduces the deep narrow furrows or fissures present on the concave sides of the real cusps (Fig 2A and 2B). In contrast, these features are lost when the enamel matrix is geometrically extrapolated from the EDJ, or when modeling with excess background nutrients (Fig 2C). These results support the role of a diffusion-limited-like process in the regulation of enamel matrix secretion, and underscore the distinct effects that the convex and concave EDJ features impose on the enamel distribution. Indeed, whereas the overall distribution of convex and concave features is conserved among molars of individual pigs, small differences in EDJ shape correspond to large differences in enamel features (Fig 3).

**Fig 3.**
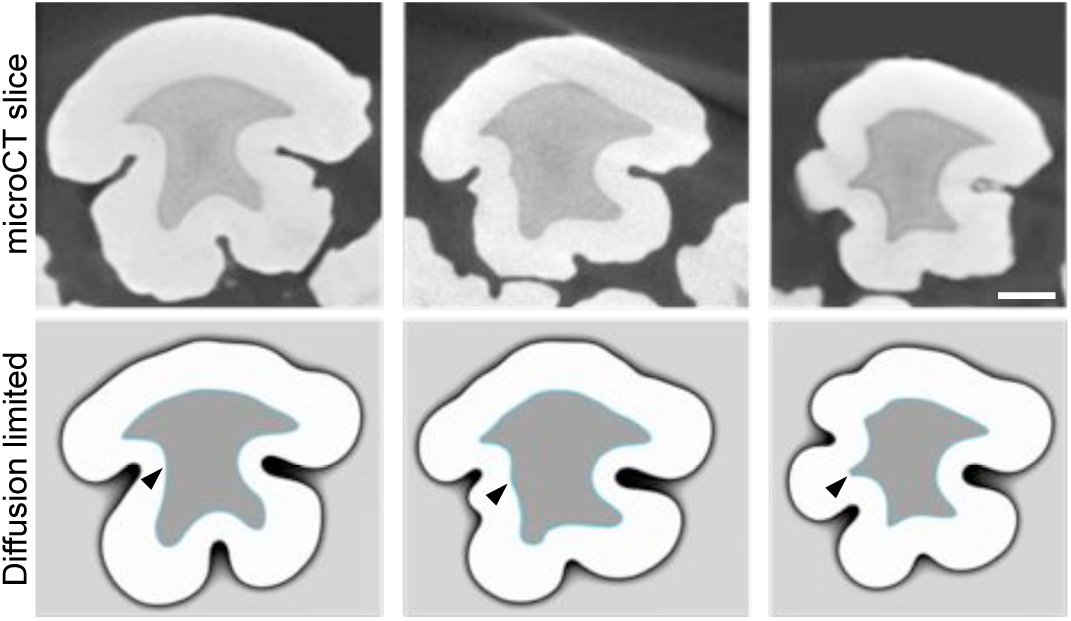
Small differences in EDJ shape produce large differences in diffusion-limited simulations. Horizontal slices of three different entoconid cusps of pig first lower molars and their diffusion-limited simulations using the same parameter values. Small differences in the size of EDJ ridges (arrowheads) correspond to large differences in enamel surface shape both in empirical and simulated enamel surfaces. The cusp in the center is 3D-simulated with the same parameters in Fig 4. Scale bar, 1 mm.

Next we simulated the matrix secretion in a whole cusp with both convex and concave features (Methods). A 3D reconstruction of these simulations show that the diffusion-limited model captures the overall enamel thickness patterns in which concavities show reduced enamel thickness whereas ridges show increased enamel thickness (Fig 4 and S1 Fig). A small ridge present in the middle of an EDJ concavity results in a local thickening of the enamel within an otherwise deep fissure (arrow heads in Fig 4), a feature completely lost in geometrically extrapolated surfaces (Fig 4D).

**Fig 4.**
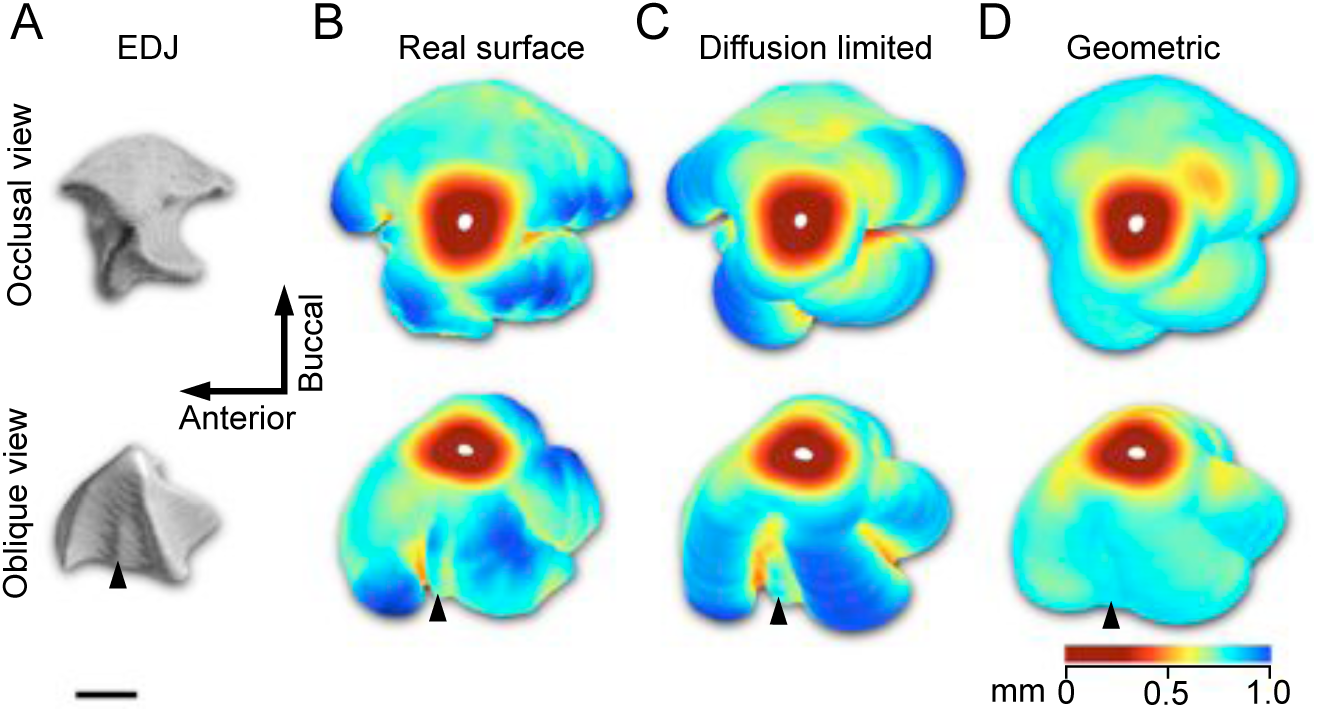
3D distribution of enamel is predicted by the diffusion-limited model. (A) The EDJ of the pig entoconid cusp (arrowhead in Fig 2B) has four ridges and only the buccal slope is convex whereas the other slopes are concave. (B) The enamel surface is thickest around the ridges and the EDJ concavities correspond to deep fissures with thinner enamel. The enamel cap, which has no EDJ, has been removed from the 3D data. (C) The diffusion-limited matrix simulation, matched to have the enamel thickness of the convex lateral slope, captures the enamel distribution patterns of the pig cusp (mean distance to the real surface is 61 μm, maximum difference = 362 μm, s.d. = 66.4). The small ridge present in the mesial slope of the cusp (arrowhead in the oblique views) corresponds to the narrow ridge of the empirical enamel surface (4B) and small ridge in an EDJ concavity (4A). (D) Geometric extrapolation from the EDJ results in relatively uniform 3D matrix thickness (mean distance to the real surface is 80 μm, maximum difference = 467 μm, s.d. = 92.0). The diffusion-limited simulations were done from 51 individual EDJ slices using the same parameter values (S1 Table). The thinner enamel in the lower parts of the lingual cusp ridge of the real cusp (towards the bottom in occlusal view) is due to the vicinity of the adjacent hypoconid cusp. Scale bar, 1 mm.

### Diffusion-limited simulations reproduce the progression of matrix secretion

In addition to the distribution of enamel in fully formed teeth, the diffusion-limited simulations can be used to examine the progression of the matrix secretion process itself. The successive positions of the matrix-secreting front during development is recorded in teeth by incremental lines (laminations or striae of Retzius, [9]) that are broadly analogous to growth rings in trees [21]. These are preserved in mature enamel and can be observed from thin sections or through phase contrast synchrotron imaging [22]. We obtained synchrotron data from a pig molar and compared the positions of individual incremental lines in convex cusp ridges with the lines in cusp concavities (Fig 5A). Both the virtual incremental lines of diffusion-limited simulations and empirical incremental lines show initially relatively uniform distances from the EDJ, but this uniformity disappears and the differences between ridges and valleys become progressively larger as the secretion accelerates in the ridges and slows down in the valleys (Fig 5). These results suggest that in addition to the final patterns of enamel distribution, the diffusion-limited model captures aspects of the actual secretion process.

**Fig 5.**
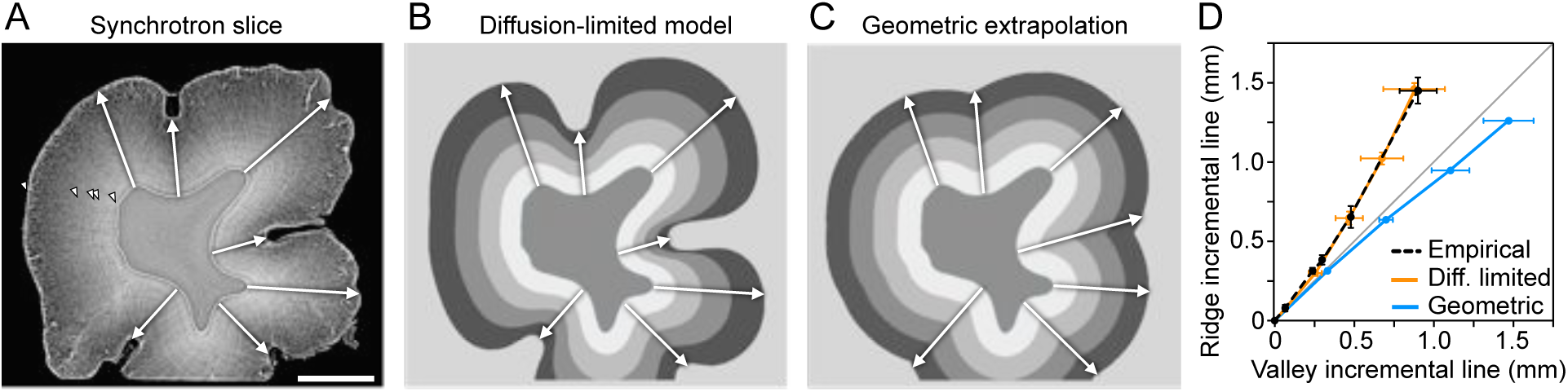
Incremental lines in pig molar tooth show comparable patterns to diffusion-limited matrix lines. (A) Incremental lines (arrowheads) are visible in a synchrotron imaged fully formed but unerupted cusp of the second lower molar. (B) Diffusion-limited simulation and (C) geometric extrapolation of matrix show contrasting patterns. The different shades of grey correspond to progressive steps in simulations (every fourth) and geometric extrapolation. (D) The arrows in Fig 5A, 5B, and 5C) show the lines that are used to measure the progression of matrix secretion in the valleys (concave EDJ regions) and ridges (convex EDJ regions). Both the empirical and diffusion-limited simulations produce progressively thicker matrix in the ridges relative to the valleys (mean lines shown, error bars denote s.d.). The border size (0.79) was set to produce the empirical enamel thickness using the same number of iterations as in Fig 3 and 4. The target enamel thickness for the simulations was measured from the left side of the cusp. Because the cusp was physically trimmed for synchrotron imaging, the section is missing the lower side. Scale bar, 1 mm.

### Subtle EDJ concavities are sufficient to produce complex enamel surface features

Domestic pigs are an example of species with relatively pointed molar cusps, allowing the simulation of matrix secretion in the horizontal plane. In contrast, molars of primates with thick enamel, including humans, typically have relatively low cusp relief. Therefore, to capture matrix secretion of human molar morphology, we run the diffusion-limited simulations vertically. Because the enamel secretion period is shorter towards the base of the tooth and the enamel is therefore thinner, we implemented a nutrient sink at the base to simulate the shorter secretion period (Methods). The sink decreases the rate of matrix formation towards the base of the crown (Fig 6A), thereby approximating crown formation and the apical decline in matrix secretion before the initiation of root development [9]. Biologically, the sink implementation can be considered a simplification of the crown-tip-to-base growth of the capillaries providing nutrients [18]. The simulations show the subtle waviness of the human EDJ, with mainly concave ripple-like features, is enough to produce the characteristic undulations of the enamel surface (Fig 6B and 6C). These features are further refined by simulations using lateral braces mimicking the presence of adjacent teeth and alveolar bone (Fig 6B), in agreement with the suggested role of the surrounding tissues in the regulation of tooth shape and mineralization [23, 24]. Taken together, these simulations indicate that even subtle EDJ features present in many hominids are important for the functional surface morphology of the tooth.

**Fig 6.**
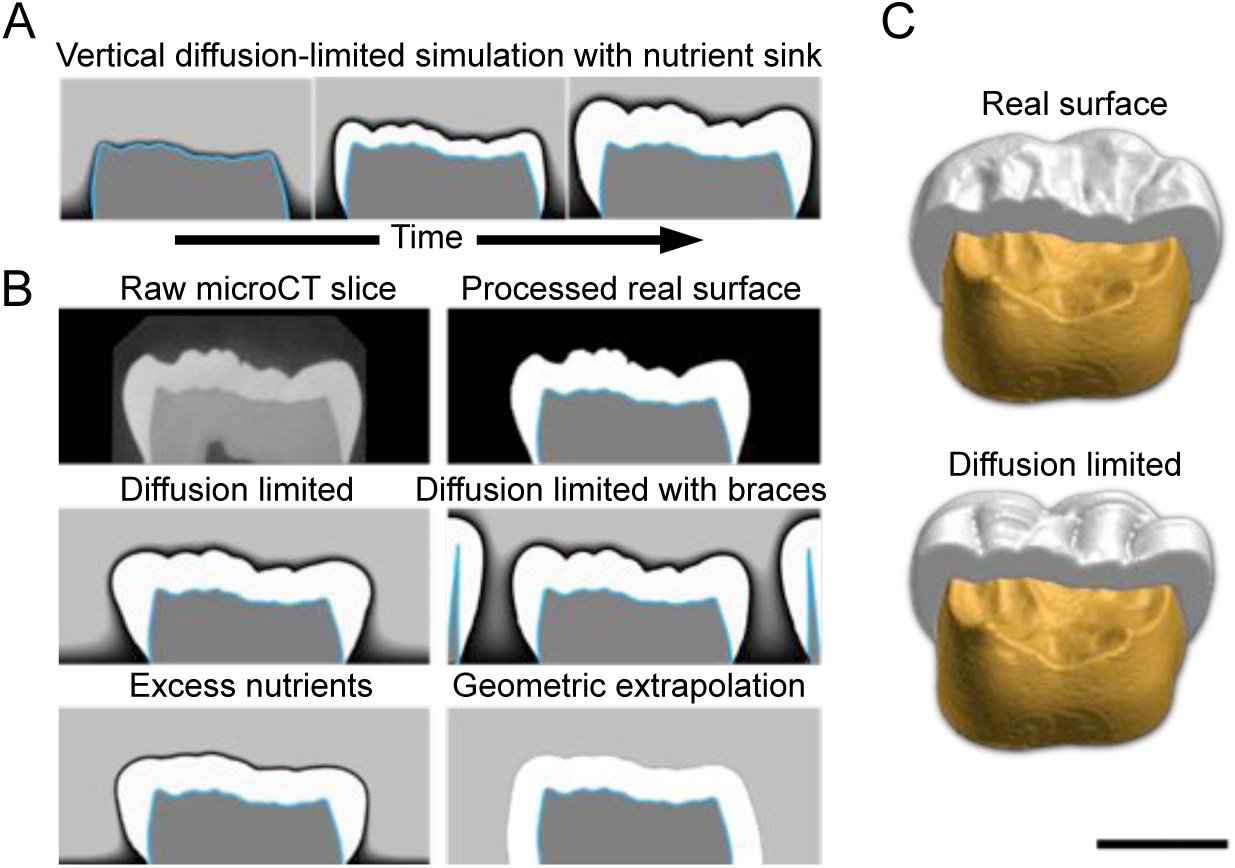
The subtle EDJ topography of human molars is sufficient to produce enamel surface features in diffusion-limited simulations. (A) To simulate the thick enamel of human molars that also have a low cusp relief, we combined vertically oriented simulations with a basal nutrient sink. The sink is used to simulate the shorter time of matrix secretion in the lower parts of the crown. (B) Diffusion-limited simulations reproduce the surface features that are lost in excess nutrient simulations and geometric extrapolations. Braces mimicking adjacent teeth and bone constrain lateral expansion of enamel matrix. (C) Obliquely lingual views of empirical and simulated enamel surfaces with a half of the EDJ (yellow) visible. The diffusion-limited simulations were done from 27 individual EDJ slices using the same parameter values (S1 Table). The tooth shown is a human third lower molar. Scale bar, 5 mm.

### Diffusion-limited matrix secretion increases surface complexity

The hallmark of diffusion-limited aggregation, which is closely related to the present model, is the fractal geometry, or self-similarity of the outcome [e.g. 25]. Although both the interfacial tension approximating the stiffness of the ameloblast layer and the background nutrient production weaken the fractal-like outcomes in our model, it is still interesting to explore the fractal nature of the enamel distribution, as it provides a quantitative measure of surface complexity. We contrasted complexities of the contours [26] of human and orangutan molars, the latter being a species with highly complex enamel distribution (Fig 7A) [14]. Box-counting dimensions (*D*_B_) of the human and orangutan molars show that although the enamel surfaces do not exhibit self-similarity per se, simulations with low background nutrient levels show stronger diffusion-limited effects (Fig 7B, S2 Table). The main factor determining enamel surface complexity of the orangutan molar appears to be the diffusion-limited effect of matrix secretion (Fig 7B). Swapping the parameter values of the human and orangutan simulations (S1 Table) produces orangutan-like enamel on human EDJ and human-like enamel on orangutan EDJ (Fig 7B, S2 Table). In contrast, excess background nutrient production results in no increase in enamel complexity relative to the EDJ in either of the species (S2 Table), indicating that diffusion-limited matrix secretion is required for the increase in surface complexity.

**Fig 7.**
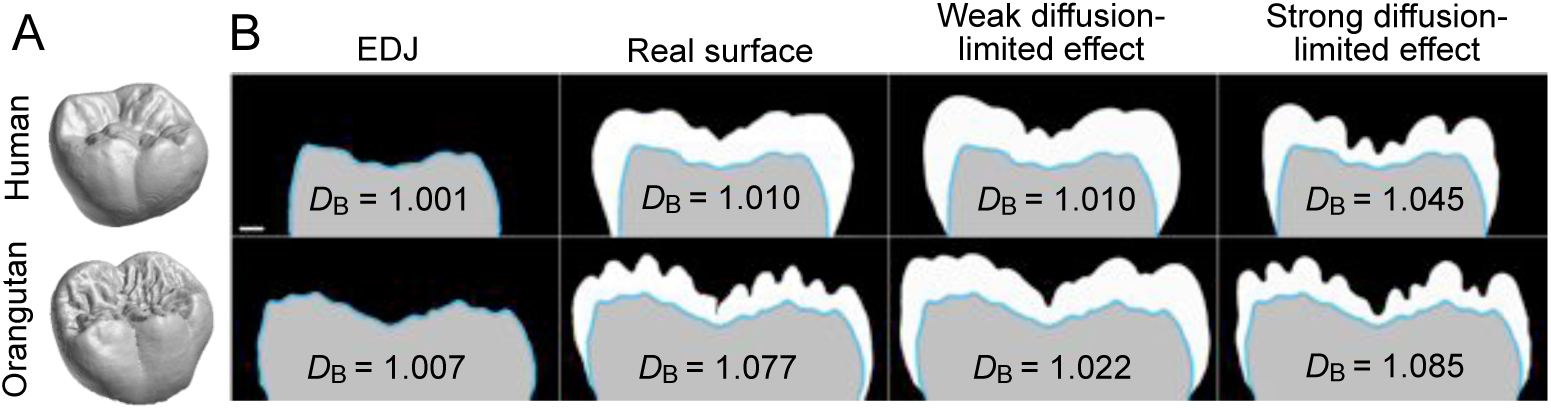
Strong diffusion-limited effect generates complex enamel surfaces. (A) Compared to human molars, orangutan molars have a large number of small surface features. (B) Vertically oriented simulations of human and orangutan molars with corresponding box counting dimensions (mean values, see S2 Table) show that lower background nutrient production, resulting in a stronger diffusion-limited effect, is enough to produce the relatively complex enamel surface of the orangutan, and increase the complexity of the simulated human enamel surface. In contrast, using human parameter values produces human-like enamel on orangutan EDJ. The human parameters are the same as in Fig 6 (S1 Table) and the longitudinal sections simulated are from approximately corresponding positions between the species. Teeth in (A) are not to scale. Scale bar, 1 mm in (B).

## Discussion

Our diffusion-limited model suggests an objective basis for mapping the EDJ morphology to enamel surface morphology. The simulations point to the critical role of the diffusion-limited effect of matrix secretion in increasing tooth surface complexity and, together with convex and concave EDJ features, in fine-tuning the shape of the enamel surface. Previously the rate of matrix secretion has been reported to be highest in the outer parts of the enamel [9, 27]. The result is supported by our simulations and synchrotron data, but only in the convex parts of the crown (Fig 4). Nevertheless, complex patterns of enamel thickness on a tooth crown can result from a single developmental process, without the need to evoke specific control or explanations for the thickness of individual features. Combined with in-depth analyses of enamel formation rates [4, 9, 27] and isotope compositions [28, 29] obtainable from sections, our model should help to move studies using enamel thickness towards more mechanistic and predictive science. The model also points to the need to investigate the role of extracellular space and capillaries in regulating diffusion, and to the potential effects of cell-cell tension in the ameloblast layer [30]. Our approach should also be applicable to other systems with extracellular matrix secretion, or organs in which directional tissue growth may be limited by a diffusion process. Diffusion-limited free boundary problems have a long history in mathematics [16, 17, 31], and as shown here, they can contribute to solving biological problems.

## Materials and methods

### Diffusion-limited model

We solve the model using a finite element method algorithm presented in [32]. The model is motivated by the classical Stefan problem [16, 17], and our formulation follows the main principles of the problem with some modifications. The model does not consider individual cells in the system, rather it treats both the enamel matrix and the dentin as continuums.

Fundamentally, we are solely focused on how a 1-manifold interface representing the ameloblast front on a cross-sectional plane evolves over time subject to interfacial tension and restricted flow of nutrients. For mathematical and implementation details, see S1 Appendix. The Matlab source code is available at https://github.com/tjhakkin/biomatrix.

### Dental data and processing

Pig and human molar samples are described in [20] and the orangutan sample is from the Finnish Museum of Natural History (UN2720). The microCT voxel resolutions were 10 to 24 μm (pig), 17 μm (human), and 32 μm (orangutan). For the analyses, the scans were downsampled to 44 μm (pig molars) and 33 μm (human and orangutan molars). Synchrotron data was collected at beamline ID19 of the European Synchrotron Radiation Facility, with voxel resolution 2.24 μm, keV = 91, and 6000 projections in 4x accumulation mode. Synchrotron data reconstruction used Paganin style single propagation distance phase contrast. All the image processing steps after primary tomography reconstructions were carried out with Fiji 2.0 [33]. To digitize EDJs for simulations, EDJs in each individual section were traced with a freehand selection or wand tool, and the area was converted to a line (Edit/Selection/Area to Line). The line was then interpolated with an interval of 10 pixels (Edit/Selection/Interpolate), fitted to a spline (Edit/Selection/Fit spline), and adjusted manually to follow the EDJ if needed. The splines were saved in ROI Manager (Analyze/Tools/ROI manager/more/save). The splines were saved as XY coordinates (File/Save As/XY Coordinates) that were converted to level sets using a Python script, included with the source code. During the conversion the spatial node density and the relative size of the EDJ within the domain were also defined. To scale different sized EDJs uniformly (e.g., when EDJs become smaller towards the cusp tip), two small triangles placed in diagonally opposing corners were included in each level set conversion. To enhance the visualization of the incremental lines in synchrotron reconstructions, three adjacent slices were averaged.

### Simulations and analyses

The main simulation parameters are listed in S1 Table. Simulation output for each step is an image file. The pig trigonids and talonids were simulated separately (Fig 2B). To compensate for the isolated entoconid cusp being larger (Fig 3) than when part of the talonid (Fig 2), interfacial tension and number of iterations were decreased in the individual cusp simulations. For simulations of multiple sections, all the sections of the analyzed step were merged into a stack in Fiji. For the pig molar cusp (Fig 4), every second microCT slice (20 μm interval, 51 slices), and for the human molar (Fig 6C), every fifth slice (66 μm interval, 27 slices) was simulated. We used the basal sink to approximate developmental progression in the vertical simulations. Because intercuspal regions lack the sink, vertical simulations slightly exaggerate enamel thickness in valleys relative to cusp tips. Geometric extrapolation of enamel thickness was obtained in Fiji by fitting a fixed sized circle along the EDJ (Process/Morphology/Gray Morphology/Dilate). To visualize enamel thickness in 3D, the EDJ and enamel surfaces were exported from Fiji (surfaces exported as Wavefront.obj from 3D Viewer plugin) and imported into Meshlab (http://meshlab.sourceforge.net/). In Meshlab, Hausdorff distance was used to compare distances between two surfaces [34]. The distances were calculated after smoothing the meshes with Laplacian smooth (3 steps). All 3D visualizations use orthographic projections.

We calculated box-counting dimensions from microCT slices of human and orangutan molars. Binary contours were generated after thresholding in Fiji (Process/Binary/Outline) and analyzed using Fiji FracLac 2.5 plugin [26]. Twelve different starting positions for the grids were used, and the contours were rotated at 36 degree steps (resulting in total of 120 *D*_B_ values for each contour). Default sampling size was used for the scaling method. We analyzed the contours with (S2 Table) and without the outer walls of the profiles, and the pattern of results remained the same.

## Supporting information

**S1 Fig. Enamel area and surface perimeter in horizontal slices of a pig cusp.** Whereas both the diffusion-limited simulation and geometric extrapolation of Fig. 4 approximate the amount of real enamel, only the diffusion-limited simulation reproduces the length of the perimeter of the real surface. The drop in enamel perimeter in the diffusion-limited simulation towards the cusp tip relative to empirical data is due to the horizontal simulations not capturing the relatively round EDJ tip. (PDF)

**S1 Table. EDJ conversion and simulation parameters.** Node density denotes the number of nodes along the longest axes in the regular triangular mesh. ‘Border’ is the proportional size of the margins surrounding the EDJ shape. ‘Base’ sets the y-axis position of the base in simulations using a sink. All EDJ simulations are run with Neumann domain boundaries, except for the simulations using basal sink for which we use mixed boundaries (Dirichlet/Neumann), with a fixed time step 0.001. All parameters are dimensionless and their absolute values have no meaning. (PDF)

**S2 Table. Box-counting dimensions calculated for human and orangutan molar teeth.** The *D*_B_ values are calculated for the binary outlines of the EDJs and enamel surfaces. The background nutrient production values for the excess background nutrient, weak and strong diffusion-limited effect simulations are 160, 75, and 30, respectively. (PDF)

**S1 Appendix. A detailed implementation of the diffusion-limited model.** (PDF)

### Acknowledgements

We thank H. Suhonen for help with microCT imaging, and J. Laakkonen for help with material. We acknowledge the European Synchrotron Radiation Facility for provision of synchrotron radiation facilities and thank P. Tafforeau for assistance in using beamline ID19. We thank M. Fortelius and the members of the Center of Excellence in Experimental and Computational Developmental Biology Research for discussions or advise.

## Author Contributions

**Conceptualization:** Teemu J. Häkkinen, S. Susanna Sova, Jukka Jernvall.

**Data curation:** Teemu J. Häkkinen, S. Susanna Sova, Jukka Jernvall.

**Funding acquisition:** Teemu J. Häkkinen, Jukka Jernvall.

**Investigation:** Teemu J. Häkkinen, S. Susanna Sova, Ian J. Corfe, Jukka Jernvall.

**Methodology:** Teemu J. Häkkinen, S. Susanna Sova, Antti Hannukainen, Jukka Jernvall.

**Project administration:** Jukka Jernvall.

**Resources:** S. Susanna Sova, Ian J. Corfe, Leo Tjäderhane, Jukka Jernvall.

**Software:** Teemu J. Häkkinen.

**Supervision:** Leo Tjäderhane, Antti Hannukainen, Jukka Jernvall.

**Validation:** Teemu J. Häkkinen, S. Susanna Sova, Jukka Jernvall.

**Visualization:** Teemu J. Häkkinen, S. Susanna Sova, Jukka Jernvall.

**Writing – original draft:** Teemu J. Häkkinen, S. Susanna Sova, Jukka Jernvall.

**Writing – review & editing:** Teemu J. Häkkinen, S. Susanna Sova, Ian J. Corfe, Leo Tjäderhane, Antti Hannukainen, Jukka Jernvall.

